# RNA Gelation in Repeat Expansion Disorders

**DOI:** 10.1101/100719

**Authors:** Ankur Jain, Ronald D. Vale

## Abstract

Expansions of short nucleotide repeats in the protein coding and non-coding regions of >30 genes produce a variety of neurological and neuromuscular disorders including Huntington’s disease (CAG repeats), muscular dystrophy (CTG repeats) and amyotrophic lateral sclerosis (GGGGCC repeats) [1-3]. Expression of expanded repeats alone is sufficient to recapitulate disease pathology in animal models [4-6]. Repeat-containing transcripts accumulate in the nucleus as aberrant “RNA foci” [7-10] and sequester numerous RNA binding proteins [11,12], leading to a disruption of cellular homeostasis [13,14]. Interestingly, RNA foci, as well as the disease symptoms, only manifest at a critical threshold of nucleotide repeats: >30 for CAG/CTG expansions [1] and >7 for the GGGGCC expansion [15]. However, the reason for this characteristic threshold, as well as the molecular mechanism of foci formation, remain unresolved [16]. Here, we show that nucleotide repeat expansions in RNA create templates for multivalent Watson-Crick (CAG/CUG expansions) or Hoogsteen (GGGGCC expansion) base-pairing. These multivalent interactions cause purified RNAs containing repeat expansions to undergo a sol-gel transition and form micron-sized clusters. Reflecting an increase in the valency for intermolecular hybridization, the gelation of purified RNA only occurs above a critical number of trinucleotide or hexanucleotide repeats. These thresholds for *in vitro* RNA gelation are similar to those associated with manifestation of disease. By visualizing RNA in live cells, we show that nuclear foci form as a result of phase separation of the repeat-containing RNA and that these foci can be dissolved by agents that disrupt RNA gelation *in vitro*. Analogous to protein aggregation disorders, our results suggest that the sequence-specific gelation of RNAs could be a contributing factor to neurological disease.

---

The RNAs associated with most repeat expansion diseases contain a regular pattern of Gs and Cs and can form intramolecular hairpins [17]. Because of their repeating nature, these sequences can also, in principle, template multivalent intermolecular interactions (Fig. 1a). Such multivalent interactions could potentially lead to the formation of large clusters via liquid-liquid phase separation [18,19] or a sol-gel transition [20-22]. To examine whether repeat-containing RNAs assemble into large clusters, we synthesized fluorescently-labeled RNAs containing 47 triplet repeats of CAG (47xCAG), which is the expansion underlying Huntington’s disease and spinocerebellar ataxias or CUG (47xCUG), which produces myotonic dystrophy and HDL2 [1]. As controls, we used RNAs of equivalent length but with arbitrary sequences with 30-75% GC content and RNAs with identical base composition as 47xCAG and 47xCUG RNAs but with scrambled sequences. Upon annealing (see Methods), the 47xCAG and 47xCUG repeat-containing RNAs formed micron-sized spherical clusters, while the control RNAs remained soluble (~5 μM for each RNA) (Fig. 1b, Supplementary Fig. 1a, 1b). The spherical clusters were >100-fold enriched in the RNA as compared to the solution phase and contained nearly one half of the RNA in the reaction (Supplementary Fig. 1c). As expected, these clusters were dissolved by RNaseA, but not proteinase K or DNase I (Supplementary Fig. 1d), confirming that the RNA clustering is not mediated by protein or DNA contaminants.

**Figure 1.**
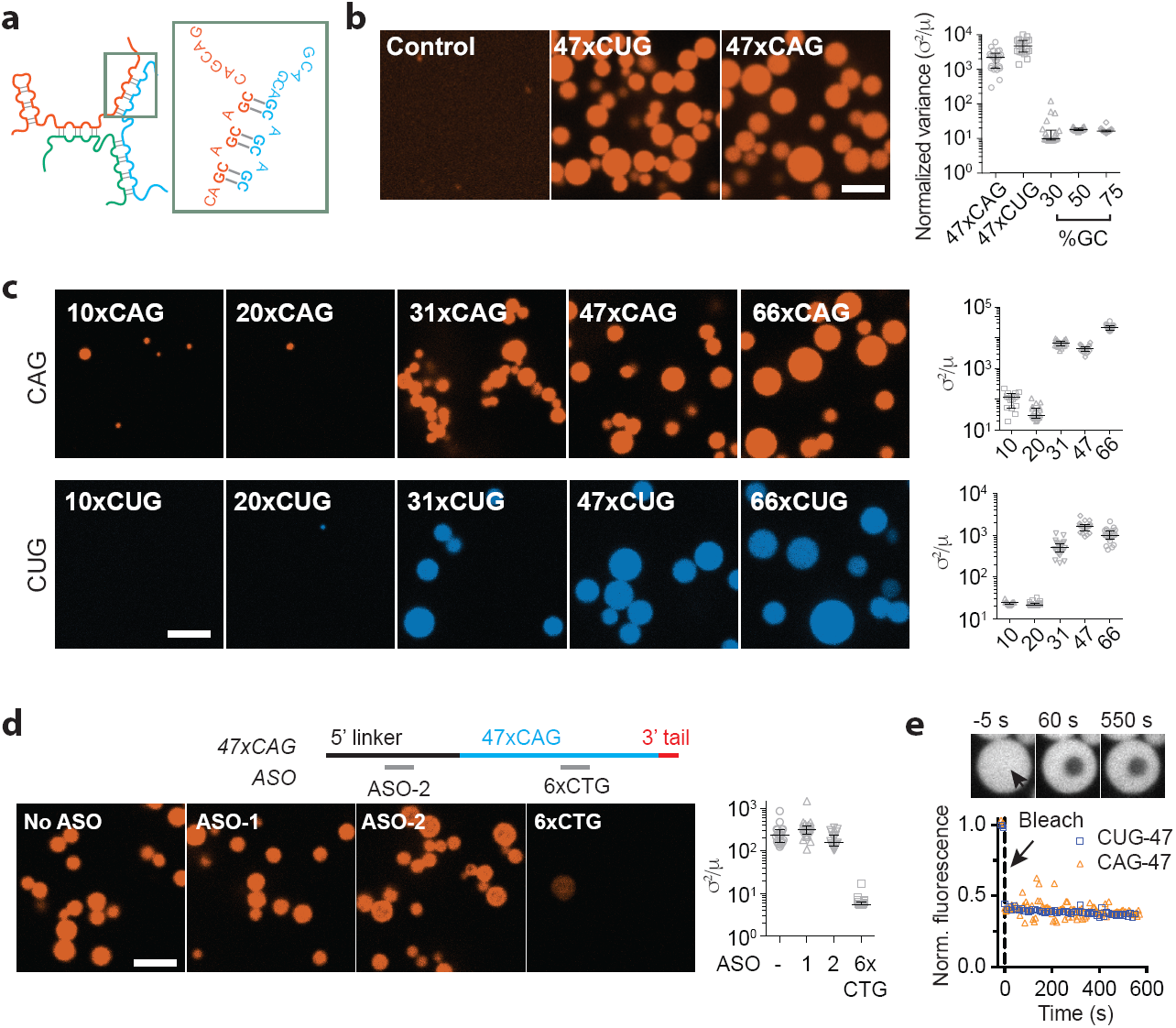
Triplet repeat containing RNAs undergo gelation *in vitro* at a critical number of repeats. (a) Schematic for possible multivalent base-pairing and multimerization of CAG repeat RNA. (b) Representative fluorescence micrographs for a control RNA with 75% GC content (left), 47xCUG (center) and 47xCAG (right) RNA. Quantitation of the inhomogeneity in the sample as normalized variance (σ^2^/μ); higher values reflect higher inhomogeneity (see Methods); each data point represents an independent imaging area (1800 μm^2^). (c) Representative micrographs of RNA with different CAG- and CUG- repeat number (200 ng/μl RNA concentration). Data are quantified as in (b). (d) DNA oligonucleotides are designed against the 47xCAG RNA that can hybridize with the CAG-repeats (6xCTG), with the 5’ linker sequence (ASO-2) or without sequence complementarity (ASO-1). 6xCTG blocks clustering of 47xCAG RNA, whereas the control oligonucleotides, ASO-1 and ASO-2 do not. (ASO: antisense oligonucleotide) (e) Fluorescence time-trajectories for 47xCUG and 47xCAG RNA clusters indicating that the clusters do not exhibit fluorescence recovery upon photobleaching. Sample images for 47xCAG RNA (top) at indicated time points before and after photobleaching. Dashed line depicts the time point of photobleaching. Scale bars are 5 μm. Error bars depict median and interquartile range. Data are representative of n = 3 or more independent experiments.

Molecules that form multivalent interactions show abrupt phase transitions with increasing valency of interaction [18-20]. An increase in the number of triplet repeats results in an increased valency for intermolecular hybridization. Consistent with this valency dependence, we find that the formation of spherical CUG and CAG RNA clusters occurred only with 30 or more triplet repeats (Fig. 1c). The intermolecular interactions between the transcripts could potentially be competed by shorter complementary antisense oligonucleotides. Indeed, addition of a 6xCTG antisense oligonucleotide prevented clustering of 47xCAG RNA, while control oligonucleotides that either cannot hybridize with the RNA or bind outside the repeat region did not (Fig. 1d). Collectively, these experiments indicate that intermolecular base-pairing interactions in the CAG/CUG-repeat region can lead to the assembly of these disease-associated RNAs in to micron sized clusters.

We next analyzed the physical properties of the CAG/CUG RNA clusters. Their spherical shapes (aspect ratio 1.05 ± 0.1, mean ± s.d., n = 214) are characteristic of polymers undergoing liquid-liquid phase separation [23]. Molecules within the liquid phase are mobile and undergo fast internal rearrangement. However, fluorescence photobleaching of the RNA in the clusters revealed little or no fluorescence recovery over ~10 min, indicating that the RNA in the clusters is immobile (Fig. 1e, Supplementary Fig. 2a, b, Supplementary Movie 1) and thus in a solid-like state. Because of their solid-like behavior, we refer to these CAG/CUG RNA clusters as “RNA gels” [21,22,24]. We hypothesize that these RNAs initially phase separate into spherical liquid-like droplets but then rapidly become cross-linked into a gel due to increasing intermolecular base-pairing. Consistent with this idea, we occasionally (in ~ 1% of the clusters) find evidence of incomplete fusion events where two droplets solidified prior to complete relaxation to a single spherical geometry (Supplementary Fig. 2c). As further evidence for this model, we found that single stranded DNA (polyT) in the presence of polyvalent cations formed liquid-like droplets but that the incorporation of multivalent base-pairing sites imparted solid-like properties to the DNA droplets (Supplementary Fig. 3, Supplementary Movie 2). Similar liquid-to-solid phase transitions have also been observed in proteins with increasing strength of intermolecular interactions [25,26]. In summary, multivalent base-pairing interactions lead to the gelation of CAG/CUG triplet repeat containing RNA *in vitro* at a similar critical number of repeats as observed for the manifestation of clinical symptoms.

Using in situ hybridization on tissues from patient biopsies, several groups have described the accumulation of trinucleotide repeat-containing RNA in aberrant nuclear foci [8]. We wished to establish a live cell reporter assay to follow the repeat-containing RNAs after transcription and determine if they assemble into nuclear foci. To enable live cell RNA visualization, we tagged the repeat-containing RNAs with 12x MS2-hairpin loops [27] and co-expressed YFP-tagged MS2-coat binding protein (Fig. 2a). Multiple stop codons upstream of the repeats were included to minimize translation of repeat containing proteins [28] (Fig. 2a). After induction of 47xCAG (Fig. 2b, Supplementary Fig. 4a) or 120xCAG (Supplementary Fig. 4b) RNA transcription in U-2OS cells, numerous nuclear foci appeared as early as 1 h post induction (Supplementary Fig. 4c, Supplementary Fig. 5, Supplementary Movie 3). The RNA foci recruited endogenous Muscleblind-like-1 protein (Supplementary Fig. 4d), sequestration of which has been previously implicated in CAG/CUG RNA toxicity [5,11]. The number of foci per nucleus increased with higher levels of RNA induction (Supplementary Fig. 4e) and with increasing repeat number (Fig. 2c). In contrast, 5xCAG (Fig. 2b, c) or control RNAs with coding or non-coding sequences (Supplementary Fig. 4f) did not form nuclear puncta.

**Figure 2.**
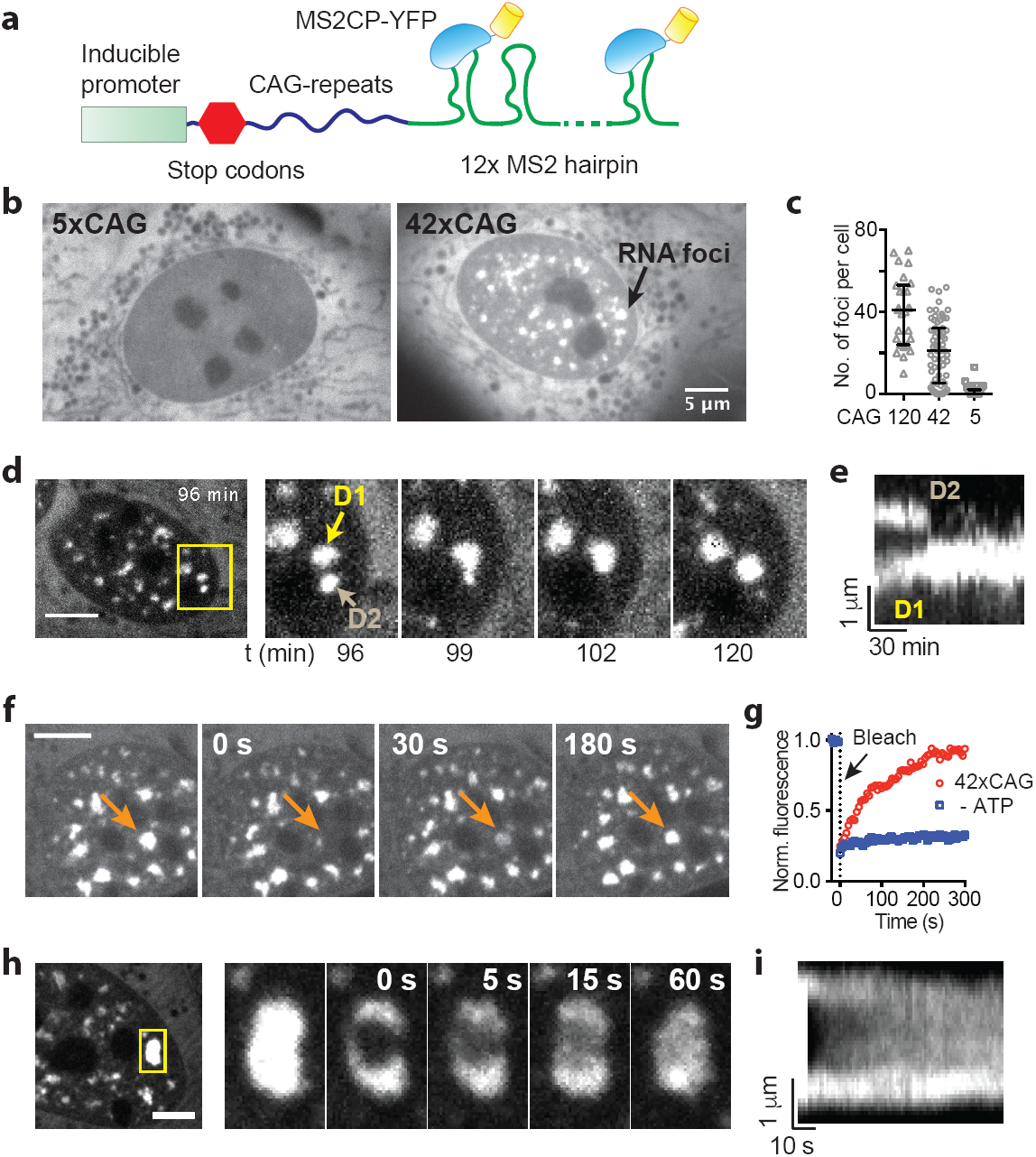
CAG repeat-containing RNAs coalesce in to liquid-like nuclear foci. (a) Schematic for RNA visualization in cultured U-2OS cells. (b) MS2-YFP fluorescence images showing accumulation of RNA as nuclear foci in cells expressing 47xCAG (right) but not 5xCAG (left). (c) Number of foci per cell as a function of CAG repeat number. Each data point represents one cell. Error bars depict median and interquartile range. (d, e) Fusion event where two adjacent RNA foci merge in to one. Images at indicated time-points after induction (d) and a corresponding kymograph (e). (f) Fluorescence micrographs for 47xCAG RNA foci before and after photobleaching at indicated time points after photobleaching. Arrow depicts site of bleaching. (g). Fluorescence recovery plots after photobleaching for 47xCAG RNA punctum before (47xCAG) and after ATP depletion (- ATP). (h, i) Partial photobleaching of a 47xCAG RNA punctum. Fluorescence micrographs at indicated time points before and after photobleaching (h) and a corresponding kymograph (i). Scale bars are 5 μm. Error bars depict median and interquartile range. Data are representative of n = 3 or more independent experiments.

Real-time microscopy revealed that the 47xCAG and 120xCAG RNA foci exhibit liquid-like properties. For example, two or more foci that came in close contact could fuse with one another (Fig. 2d, 2e, Supplementary Movies 3, 4), a hallmark of liquid-like behavior as has been previously shown for various physiological membraneless organelles [29,30]. Upon photobleaching, nuclear puncta exhibited near-complete fluorescence recovery (83 ± 13% recovery, τ_FRAP_ = 81 ± 24 s; mean ± s.d., n = 5 foci across 3 independent experiments), indicating that the RNA is dynamic and can move into and out of the nuclear foci (Fig. 2f, g, Supplementary Movie 5). Upon photobleaching a portion of a 47xCAG RNA punctum, the fluorescence also recovered rapidly (τ_FRAP_ = 18 ± 5 s, mean ± s.d., n = 5 foci) (Fig. 2h, i, Supplementary Movie 6), suggesting that RNA can undergo internal rearrangement within the foci, which is further indicative of liquid-like behavior [29]. Thus, unlike the solid-like behavior of clusters prepared from pure CAG RNA *in vitro* (Fig. 1e), CAG RNA nuclear puncta in cells display liquid-like properties. We hypothesized that the increased dynamicity might arise from specialized proteins (e.g. helicases) in the nucleoplasm that remodel RNA base-pairing. Consistent with this hypothesis, depletion of cellular ATP significantly reduced fluorescence recovery of the RNA foci upon photobleaching (23 ± 7% recovery, mean ± s.d., n = 7, Fig. 2g, Supplementary Movie 7).

Next, we asked if perturbations that prevent RNA gelation *in vitro* also affect the stability of RNA foci in cells. We found that doxorubicin, a nucleic acid intercalator that disrupts base pairing [31], blocked the formation of CAG RNA gels *in vitro* (Fig. 3a). Similarly, addition of doxorubicin (2.5 μM) potently dissolved the 47xCAG nuclear foci in cells (Fig. 3b). Second, phase separation of anionic polymers, like RNAs, is promoted by multivalent cations, which can bridge between polymers, and is inhibited by monovalent cations that compete for occupancy with the multivalent cations [32,33]. We found that the gelation of the CAG repeat-containing RNAs was inhibited by monovalent cations such as NH_4_^+^ (Fig. 3c). To test the effect of monovalent cations in cells, we used ammonium acetate, which readily permeates into cells and does not perturb intracellular pH [34]. Strikingly, the addition of 100 mM ammonium acetate led to the disappearance of 47xCAG RNA foci within minutes (Fig. 3d, Supplementary Movie 8). To preclude the possibility that ammonium acetate or doxorubicin cause the dissociation of MS2-coat binding protein from the RNA, we performed RNA fluorescence in situ hybridization experiments and observed a similar reduction in the number of nuclear foci (Supplementary Fig. 6). Thus, agents that inhibit the gelation of purified repeat-containing RNAs *in vitro* also disrupt nuclear foci in cells.

**Figure 3.**
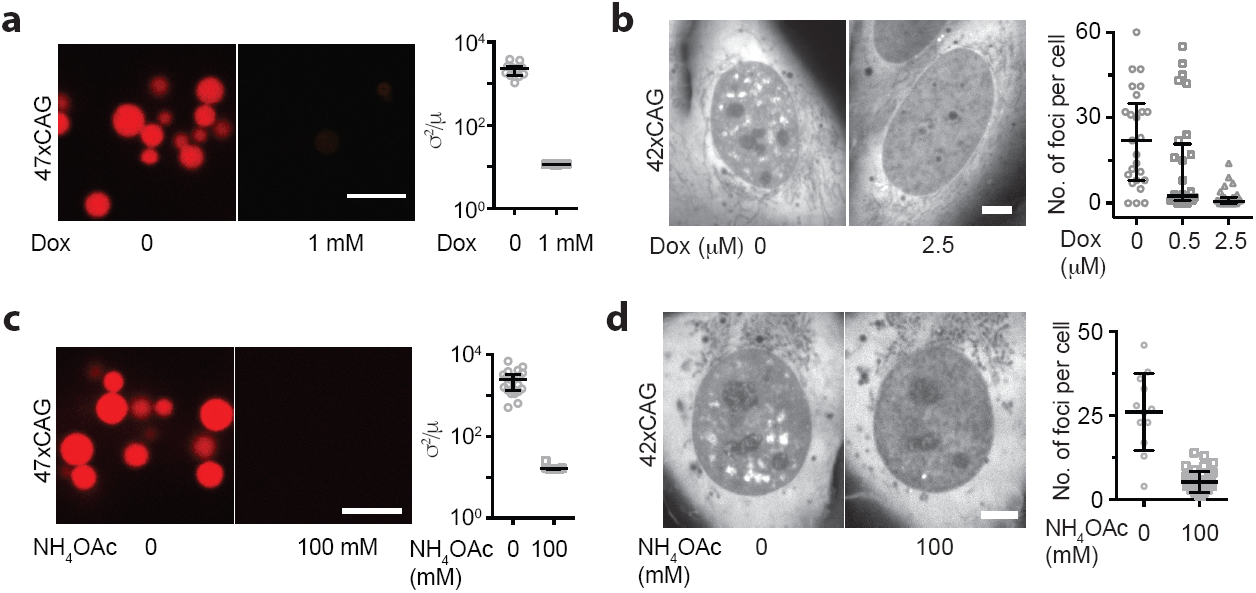
Nuclear foci are reversed by agents that disrupt RNA gelation *in vitro*. (a) Representative fluorescence micrographs and corresponding quantification of inhomogeneity for 47xCAG RNA *in vitro* with or without 1 mM doxorubicin. (b) Fluorescence micrographs of 47xCAG expressing cells with or without treatment with 2.5 μM doxorubicin. Quantification of the number of foci per cell 2 h after treatment with doxorubicin at indicated doses. (c) Same as (a) for 47xCAG RNA with or without 100 mM NH_4_OAc. (d) Treatment of 47xCAG expressing cells with 100 mM NH_4_OAc disrupted RNA foci. Images of a cell before and 300 s after treatment with NH_4_OAc. Quantification of the number of foci per cell before and 10 min after treatment with 100 mM NH_4_OAc. Scale bars are 5 μm. Error bars depict median and interquartile range.

Besides the canonical Watson-Crick base pairing, nucleic acids may also form Hoogsteen base-pairs such as in G-quadruplexes. Interestingly, RNA with the ALS/FTD-associated GGGGCC repeat expansion found in the *C9orf72* locus, the most common genetic cause of ALS [35], is shown to form G-quadruplexes *in vitro* and *in vivo* [36,37]. A single G-quartet can bring up to four RNA strands together, and the repeating G-quadruplex as found in *C9orf72* could potentially give rise to multimolecular RNA complexes [36] (Fig. 4a). Additionally, G-quadruplexes form stronger interactions than Watson-Crick base pairing [38]. Indeed, we find that an RNA with 3xGGGGCC is largely soluble but as few as 5x repeats of GGGGCC forms spherical clusters (Fig. 4b). Longer repeats (10x and 23x) resulted in the formation of an interconnected mesh-like network of aggregated RNA (Fig. 4b). In contrast, 23xCCCCGG RNA, which can form multivalent Watson-Crick base pairing but not G-quadruplexes, remained soluble under the same conditions in which 23xGGGGCC formed aggregates (Fig. 4c). As described for the CAG/CUG RNA gels, the GGGGCC RNA clusters are also solid-like and did not exhibit fluorescence recovery after photobleaching (Supplementary Fig. 7). In summary, repetitive GGGGCC RNA assemble in to supramolecular aggregates *in vitro* by forming multimolecular G-quadruplexes at a lower repeat number than repetitive CAG/CUG RNA.

**Figure 4.**
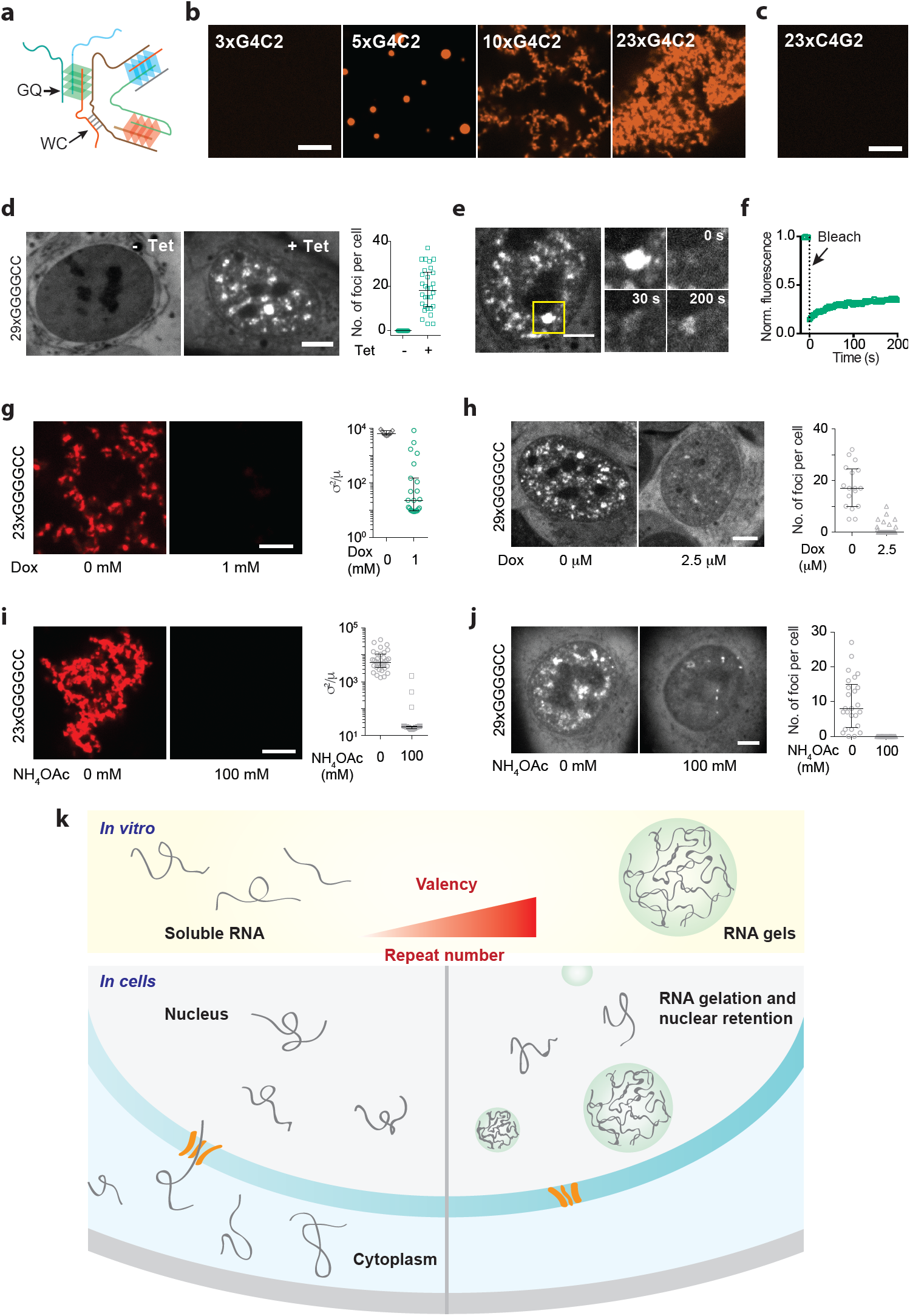
Gelation of GGGGCC repeat containing RNA and model for RNA foci formation. (a) Schematic for GGGGCC RNA multimerization (GQ: G-quadruplex, WC: Watson-Crick). (b) Representative micrographs for GGGGCC RNA at indicated number of repeats. (c) 23xCCCCGG RNA is soluble under similar conditions. (d) Expression of 29xGGGGCC RNA leads to formation of nuclear foci in U-2OS cells. (e, f) 29xGGGGCC RNA foci exhibit incomplete fluorescence recovery upon photobleaching. Micrographs at indicated time points (e) and recovery plot (f). (g) Representative fluorescence micrographs and corresponding quantification of inhomogeneity for 23xGGGGCC RNA *in vitro* with or without 1 mM doxorubicin. (h) Fluorescence micrographs of 29xGGGGCC expressing cells treated with 2.5 μM doxorubicin for 2 h (right) or DMSO only (left). Quantification of the number of foci per cell with or without doxorubicin treatment. (i) Same as (g) with or without 100 mM NH_4_OAc (j) Treatment of 29xGGGGCC RNA expressing cells with 100 mM NH_4_OAc disrupted RNA foci. Images of a cell before and 300 s after treatment with NH_4_OAc. Quantification of the number of foci per cell before and 10 min after treatment with 100 mM NH_4_OAc. Scale bars are 5 μm. Error bars depict median and interquartile range. (k) Model for RNA foci formation in repeat expansion disorders. The repeat expansion sequences form templates for intermolecular base pairing. An increase in the repeat number leads to an increased valency of intermolecular interaction, and thus an increased propensity to undergo gelation. In cells, the RNA clusters are retained in the nucleus and may disrupt cellular homeostasis.

Expression of 29xGGGGCC RNA in U-2OS cells also led to the formation of nuclear puncta (Fig. 4d). However, the GGGGCC RNA foci exhibit incomplete recovery after fluorescence photobleaching (36 ± 16% recovery, mean ± s.d., n = 5, Fig. 4e, f, Supplementary Movie 9), indicating that the GGGGCC RNA foci are less dynamic than the CAG RNA foci. This result is consistent with a stronger inter-molecular interaction between the GGGGCC quadruplex (Fig. 4a). Lastly, treatment with doxorubicin or ammonium acetate disrupted GGGGCC RNA gels *in vitro* as well as the RNA foci in cells (Fig. 4g-j) indicating that both base-pairing and electrostatic interactions are essential for the assembly of GGGGCC RNA foci.

In summary, we have shown that the propensity of an RNA to form multivalent canonical Watson-Crick (in CAG/CUG expansions) or Hoogsteen (in GGGGCC expansions) base-pairing can lead to its gelation without requiring protein components. An emerging body of research has shown that RNP granules, such as stress granules, P-granules and nucleolus, are phase separated compartments [24]. Numerous recent studies have characterized proteins with regard to their ability to phase separate and mediate the assembly of these RNP granules [26,39,40]. Our results demonstrate that sequence-specific base-pairing properties of RNAs can lead to their phase separation and gelation, and raise the possibility that such phenomena could contribute to physiological granule assembly as well.

In the case of diseases produced by nucleotide repeat expansions, our data suggest that intermolecular RNA base-pairing can result in the aggregation and sequestration of RNA into nuclear foci (Fig. 4k). RNA gelation, which occurs at a boundary condition of increasing valency, might explain why a critical threshold of nucleotide repeats must be reached to produce disease (Fig. 4k), a puzzling phenomenon that has lacked a mechanistic explanation thus far [1,16]. RNA gelation and foci formation likely occur upstream of known drivers of cellular toxicity such as sequestration of RNA binding proteins [11,12], nucleolar stress [41] and disruption of nucleocytoplasmic transport [42,43]. In addition to these direct consequences of nuclear RNA aggregation, aberrant protein production due to canonical [44] or non-ATG translation [45] from repeat expansions also likely contributes to toxicity and neurological disease. Our results may explain why placement of distinct repeat expansions in seemingly unrelated genes can result in similar clinical syndromes [46,47]. In addition, our findings explain how RNA duplex [48,49] or quadruplex [50] destabilizing compounds disrupt nuclear foci and suggest that new strategies for disrupting RNA-RNA base pairing could lead to novel therapeutics for treating repeat expansion diseases.

## Acknowledgments

We thank M. K. Rosen, W. W. Seeley and the members of Vale lab for helpful discussions. A.J. is supported by the Damon Runyon Cancer Research Foundation postdoctoral fellowship. R.V. is an investigator with the Howard Hughes Medical Institute.

